# Mutation-biased adaptation: The Yampolsky-Stoltzfus model revisited

**DOI:** 10.64898/2026.10.05.756661

**Authors:** Su-Chan Park, Joachim Krug

## Abstract

The Yampolsky-Stoltzfus (YS) model provided the first formal demonstration that a bias in the introduction of new mutations can shape the direction of adaptive evolution, even when mutation rates are small relative to selection coefficients. Despite its foundational role, the model has been characterized almost entirely by simulation and closed-form solutions for its central quantity, the probability *P*_A_ that a mutationally favored but less-fit genotype fixes ahead of a rarer, fitter alternative, have been available only in the limiting regimes of vanishing mutation supply and infinite population size. Here we derive closed-form approximations for *P*_A_ across the intermediate regime of finite population size and appreciable mutation supply that lies between these two limits. Under strong selection between the competing mutants, we combine an establishment-probability argument with clonal interference theory to obtain an explicit expression for *P*_A_ that accounts for both the ongoing input of the mutationally favored mutation and the frequency-dependent establishment probability of its competitor. Under weak selection between competing mutants, we derive a complementary expression based on the fixation of a mixed population under Kimura’s classical formula. Both expressions agree closely with direct simulation of the underlying Wright-Fisher process, recover the known limiting cases, and remain consistent with one another over a substantial intermediate range of parameters. These results are expected to provide a basis for assessing how the strength of mutation bias affects adaptation in empirical data.

## Introduction

Mutation is the ultimate source of heritable variation, but the generation of this variation is not unbiased (Agashe et al. 2023). Certain classes of nucleotide change have been observed to arise several-fold more often than others across a wide range of species (Katju and Bergthorsson 2019). Whether such biases can shape the direction of adaptive evolution, rather than being overwhelmed by selection, was addressed formally by Yampolsky and Stoltzfus (2001) using a minimal model with a two-locus, two-peak fitness landscape, which we refer to throughout as the Yampolsky-Stoltzfus (YS) model. Assuming strong selection and weak mutation (SSWM), Yampolsky and Stoltzfus (2001) derived an analytic expression for the probability that the population adapts via one route rather than the other, which shows that a bias in the mutation rates can influence evolutionary outcomes even when mutation rates are small relative to selection coefficients. This result generalizes readily within the SSWM framework, or, equivalently, under origin-fixation dynamics (McCandlish and Stoltzfus 2014), where the rate of evolution along any pathway is taken to be the product of a mutational origination rate and a fixation probability. In this way the evolutionary bias between different alternatives factorizes into an origination bias and a fixation bias (Cano et al. 2023).

Substantial empirical support for the occurrence of mutation-biased adaptation has accumulated since the introduction of the YS model (Stoltzfus 2021). Parallel adaptive substitutions across natural and laboratory systems have been observed to show an enrichment for mutationally favored changes (Stoltzfus and McCandlish 2017; Cano *et al*. 2022), and this relationship has recently been reported to hold broadly across taxa when mutational and adaptive substitution spectra are compared directly (Gitschlag *et al*. 2025). In evolution experiments with bacteria, it has been found that a tradeoff between common low-effect and rare high-effect mutations shapes the adaptive trajectories towards antibiotic resistance in populations of different size (Schenk *et al*. 2022). Very recently, Barber and Couce (2026) designed an experiment where mutator strains of *Escherichia coli* were offered two alternative trajectories to different resistance targets and mapped out the way the outcomes depend on the competition or alignment between mutation bias and selection. Not all biologists agree on the role of mutation bias in adaptation, however, and the topic is subject to ongoing debate (Svensson and Berger 2019; Gomez *et al*. 2020; Stoltzfus 2021; Svensson 2022). To some extent this reflects the different roles that *de novo* mutations and standing variation play in microbial evolution experiments and natural populations, respectively, which bias the expectations of researchers working in these fields. In this situation, it seems desirable to develop an analytic understanding of the minimal YS model in order to clarify the parameter ranges where mutation bias can dominate adaptation.

So far, the YS model has been characterized almost entirely by simulation rather than by exact theory. As Stoltzfus (2019) has noted, the model has admitted closed-form treatment only in two limits. Either mutation supply vanishes, corresponding to the SSWM/origin-fixation regime described previously, or the population size grows without bound at fixed mutation rate, which implies that a deterministic treatment applies and the fitter type predictably prevails. An approximate treatment of the stochastic model in which the fitter genotype has infinite fitness was presented by Soares *et al*. (2021), using the somewhat questionable assumption that mutants can appear only at the first generation. While the suggested formula recovers the above two limits, the authors noted that it is not valid at intermediate population sizes.

What remains open, therefore, is the intermediate regime of finite population size with appreciable mutation supply, where no closed-form expression has yet been reported for the probability that the mutationally favored outcome fixes before the selectively favored one. Closing this gap is not merely of mathematical interest. Without a tractable and easily interpretable expression for this probability, the YS model is difficult to place in correspondence with related formal frameworks such as the nearly neutral theory (Ohta 1992), and fitting it to experimental data requires simulation-based inference that scales poorly with sample size and parameter dimension (McCandlish et al. 2015). Here we derive accurate analytical results for the YS model across this previously unresolved intermediate regime. Specifically, we obtain closed-form expressions for the probability that a population fixes the mutationally favored rather than the selectively favored peak, as an explicit function of the two selection coefficients, the population size, and the mutation supply. We treat this problem in two complementary regimes: one in which selection between the two mutants is strong, and one in which it is nearly neutral. This yields two distinct closed-form expressions that agree over an intermediate range of parameters rather than being mutually exclusive, and that together cover the great majority of biologically relevant parameter space. We further verify that both expressions recover the known SSWM and infinite-population limits as special cases.

This paper is structured as follows. In Section Model, we describe the YS model in full so that the paper is self-contained. In Section Results, we derive analytic expressions for the effect of mutation bias on adaptation, treating separately the case of strong selection between the two mutants and the case in which selection is nearly neutral. In Section Discussion, we consider applications of these results and directions for future work.

## Model

The Yampolsky-Stoltzfus (YS) model (Yampolsky and Stoltzfus 2001) is a deliberately minimal scenario designed to isolate a single question. If a population can reach a fitness improvement by two alternative mutational routes, and those routes differ both in how often they arise and in how much fitness benefit they confer, then which route does the population actually take? To ask this question cleanly, the model considers a population that starts out fixed for a single wild-type genotype (WT), which is not itself optimal. Two different mutant genotypes A and B are available to WT individuals. Biologically, A and B represent two alternative single mutational steps out of the ancestral state, each opening a distinct trajectory toward higher fitness. Denoting the WT fitness by *w*_0_ and the mutant fitnesses by *w*_A_, *w*_B_, we assume the ordering

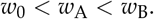

Both derived genotypes are beneficial relative to the ancestor, but B is the more beneficial of the two. This is what is meant by describing the model as a “two-locus, two-peak” fitness landscape. The ancestral genotype sits in a valley, and two separate mutational paths lead uphill to two peaks of unequal height. Because the model restricts attention to just these three genotypes and forbids any further mutation, once a population becomes monomorphic in A it can never subsequently reach the higher peak B, and vice versa. The choice between the two peaks is effectively permanent once made. This is what makes the model informative about a genuinely important biological scenario known as reciprocal sign epistasis (Poelwijk *et al*. 2007). The crossing from one adaptive solution to another is not possible without passing back through a less-fit intermediate, so that whichever beneficial mutation a population happens to fix first prohibits access to the alternative (Salverda *et al*. 2011).

The population itself is modeled as simply as possible. It is haploid and asexual, of constant size *N*, and evolving in discrete, non-overlapping generations under the standard Wright-Fisher sampling scheme (the exact mathematical rule for how offspring genotype frequencies are drawn from parental frequencies at each generation is given in Appendix A). At generation *t* = 0 every individual carries the WT genotype. The population is monomorphic at the outset, with no standing variation for either A or B. Both must be introduced by new mutation over the course of evolution. Each generation, a WT individual produces offspring that mutate independently to A with probability *µ*_A_ and to B with probability *µ*_B_. It is convenient to express the fitness advantage of each derived genotype as a selection coefficient, *s*_*i*_ = (*w*_*i*_ /*w*_0_) *−*1 for *i∈ {*A, B *}*. Throughout, the model assumes

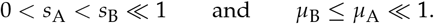

This parameterization is chosen to set up a tension between mutation and selection. Genotype A is the mutationally favored route (it arises more often) while B is the selectively favored route (it confers a larger fitness benefit). No other mutations are permitted, which guarantees that the population’s fate is decided entirely between these two alternatives. For a graphical representation of the model, see Fig. 1.

**Figure 1.**
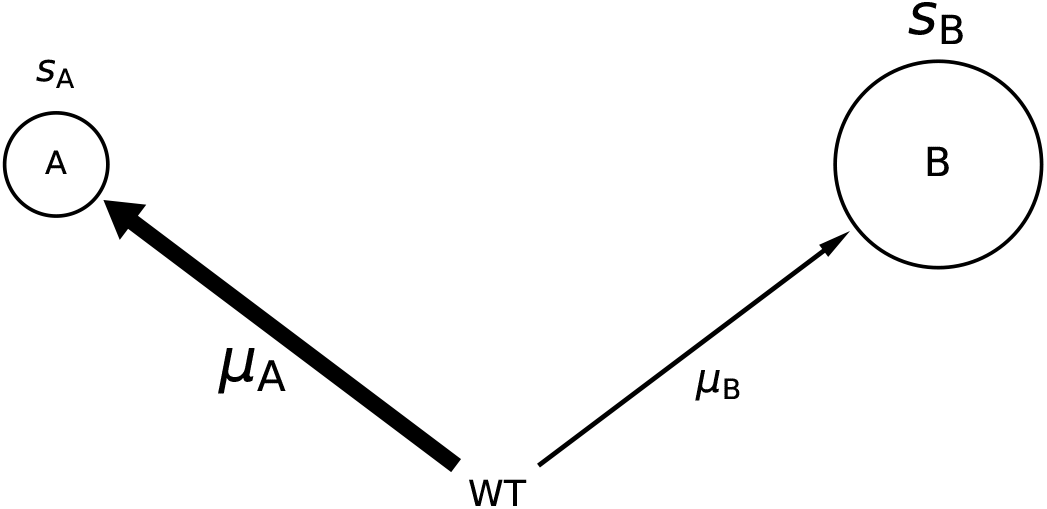
Graphical representation of the relation among the three genotypes WT, A, and B. Arrows indicate the possible mutations. Thickness of the arrows signifies the corresponding mutation rates, and the size of the circles represents the fitness of the target genotypes. In addition, larger mutation rates and selection coefficients are represented by larger font size.

The quantity of interest is the fixation probability *P*_A_ that the population ultimately fixes for the mutationally favored but less-fit genotype A rather than the more-fit but rarer B. Framed this way, *P*_A_ directly measures the extent to which a bias in the *production* of variation (through *µ*_A_ *> µ*_B_) can override a bias imposed by *selection* (through *s*_B_ *> s*_A_). If mutation were irrelevant to evolutionary direction, *P*_A_ would be small whenever *s*_B_ is appreciably larger than *s*_A_, regardless of the mutation rates.

The central result of the YS model is that this is not generally true. For a wide range of population sizes and mutation-rate ratios, *P*_A_ can be substantial, and even exceed one half, meaning that the population is more likely to fix the mutationally favored but inferior genotype than the rarer, superior one. Deriving an explicit mathematical expression for *P*_A_ in terms of *N, µ*_A_, *µ*_B_, *s*_A_, and *s*_B_ is the objective taken up in the remainder of this paper.

## Results

Since two mutant lineages can both be present in the population before either one reaches fixation, they can end up competing with one another for fixation. To obtain an analytic solution to this competition, we make use of the framework of clonal interference theory (Gerrish and Lenski 1998; Park and Krug 2007; Park et al. 2010), which describes how beneficial mutations competing for fixation in the same population interfere with one another. The basic intuition, following Jain *et al*. (2011), is simple. The weaker mutation (A) can only reach fixation if it manages to avoid having to compete directly against the stronger one (B) on its way there. Since clonal interference theory treats competing lineages as growing deterministically once they are established, it implicitly assumes that selection is strong. When selection between the two mutations is not strong, however, this assumption breaks down. Even after B has become established in the population, A can still reach fixation with non-negligible probability, an outcome that clonal interference theory does not properly capture. We therefore need a separate approach for the weak-selection case.

In what follows, we first analyze the model using clonal interference theory under the assumption of strong selection, and then turn to a separate analysis for the case of weak selection.

### Strong selection

We begin by studying the fixation probability *P*_A_ under the assumption of strong selection, i.e., in the regime where *N*(*s*_B_*−s*_A_) *≫* 1. Under this assumption, once type B has become established in the population, it is essentially guaranteed to go on to fixation, no matter how common type A has become in the meantime. In other words, A can only reach fixation in the scenario where B never manages to become established at all. This lets us write *P*_A_ as the product of two probabilities: the probability that A becomes established before B, and the probability that A subsequently goes on to fixation without interference by the stronger mutant B. We refer to this combined approach as the establishment-clonal interference (ECI) framework. As will become clear shortly, ECI includes the SSWM/origin-fixation framework discussed earlier as a special case.

By establishment of A, we mean that the lineage carrying A has grown numerous enough that it is very unlikely to be lost by chance, or genetic drift alone. As a rule of thumb, a new mutant lineage is considered established once the number of mutant copies in the population reaches the inverse of the selection coefficient (Park *et al*. 2010). We refer to the period before A reaches this point as the establishment phase.

During the establishment phase, each new mutant lineage, regardless of which genotype it carries, can be treated as evolving independently of the others. Since a lineage that has become established is very likely to fix (given that no further new mutations are allowed), the probability that a lineage becomes established can itself be approximated by its fixation probability. Under this approximation, the number of generations *t*_*i*_ that elapse before a genotype *i ∈{*A, B*}* becomes established behaves like an exponentially distributed random variable with mean

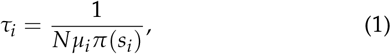

where *π*(*s*) is the probability that a single mutant copy with selection coefficient *s* eventually fixes under the assumption that no further mutation appears. In the discrete-generation Wright-Fisher model, this fixation probability is well approximated by (Kimura 1962; Park et al. 2010)

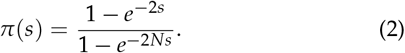

Assuming strong selection, in the sense of *Ns ≫*1, this reduces to *π*(*s*) ≈1 *− e*^*−*2*s*^, which can be further simplified to *π*(*s*) ≈ 2*s* when *s* is small.

Treating the establishment of A and B as a competition between two independent, exponentially distributed waiting times, the probability *P*_est_ that A establishes before B is

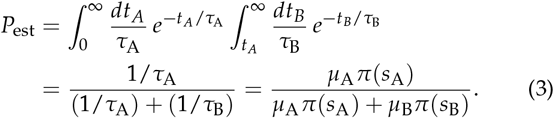

Once A has become established, and provided that no B lineage has yet appeared, the frequency *x* of mutation A in the population increases deterministically according to the differential equation (derived in Appendix A)

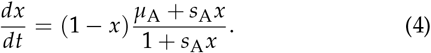

We refer to this phase of deterministic increase as the growth phase. During the growth phase, the ultimate fate of A hinges entirely on whether B subsequently manages to become established. Mutant A can fix provided B never establishes. On the other hand, if B happens to become established at any point during the growth phase, then it is thereafter highly likely to be the one that fixes instead. Figure 2 illustrates these two possible outcomes using representative simulated trajectories of genotype frequency over time. Panel (a) shows a case in which A fixes, and panel (b) a case in which B does. Panel (a) also marks, for reference, the establishment phase and the growth phase introduced above.

**Figure 2.**
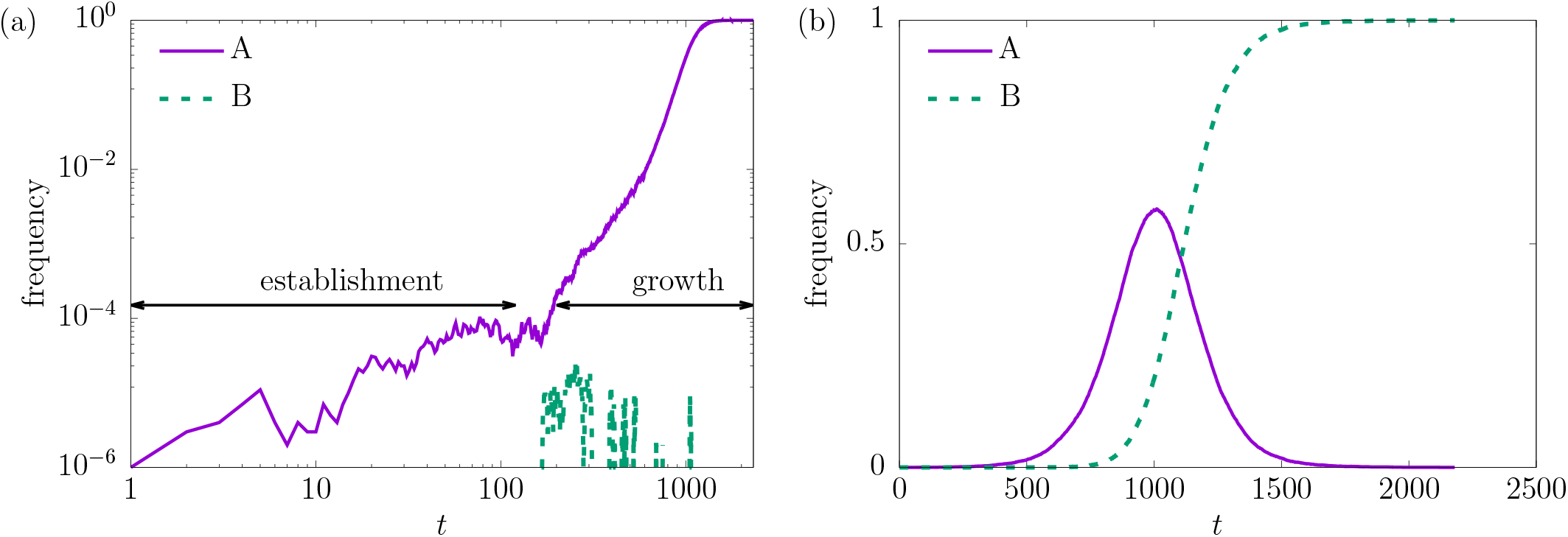
Two sample paths to fixation. Parameters used here are *N* = 10^6^, *µ*_A_ = 10^*−*6^, *µ*_B_ = 10^*−*8^, *s*_A_ = 0.01, and *s*_B_ = 0.02. Double-logarithmic plot of frequency vs generation *t* for genotype A (solid purple line) and B (dashed green line) for a realization of the process where A fixes. The logarithmic scale amplifies the early-time stochastic nature of establishment. (b) Plot of frequency vs generation *t* for a realization where B fixes. To show the deterministic nature of clonal interference clearly, we depict the frequencies on a linear scale.

Since the underlying process is Markovian (meaning that its future behavior depends only on the population’s current state, not on the specific path by which it arrived there), we can treat the establishment phase and the growth phase as statistically independent stages. The overall fixation probability *P*_A_ of A is then obtained simply by multiplying the probability *P*_est_ that A becomes established first by the probability *P*_no_ that B does not establish during the subsequent growth phase.

We now derive *P*_no_. At the point when A has reached frequency *x*, new B mutants continue to arise at rate *N*(1*− x*)*µ*_B_ (recall that only offspring of the remaining wild-type individuals can acquire the B mutation). Since at this point the population’s mean fitness relative to the WT is 1 + *s*_A_ *x*, a newly arising B mutant establishes with probability *π*(*s*_B_ *−s*_A_ *x*) rather than the naive *π*(*s*_B_). This is because its fitness advantage is effectively reduced by the fitness gain that A has already achieved. The expected number *N*_*e*_ of independent B establishment events over the course of the entire growth phase is therefore

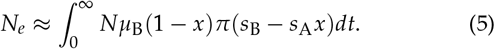

Using Eq. (4), we can change variables from *t* to *x* via

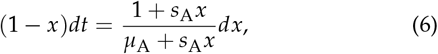

which, together with the approximation *π*(*s*) *≈* 2*s*, gives

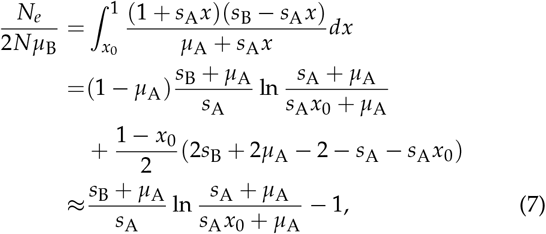

where *x*_0_ denotes the frequency of A at the start of the growth phase, that is, at the moment A first becomes established. Note that the final result of Eq. (7) is obtained also if the term *s*_A_ *x* in the numerator of Eq. (6) is neglected.

If a lineage reaches *n* copies with *n≪ N*, then the fate of every mutant can be treated independently and, accordingly, the probability that no mutant goes to fixation can be approximated by

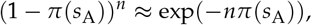

where *π*(*s*_A_) is assumed small. Thus, if *n≈* 1/*π*(*s*_A_), we can regard A as being effectively established, and we therefore set

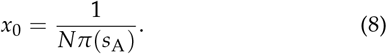

Treating the establishment of B lineages as an independent Poisson process (Gerrish and Lenski 1998), that is, as random events occurring independently of one another at a roughly constant underlying rate, the probability *P*_no_ that no such establishment event occurs at all is simply

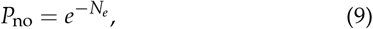

which, combined with Eq. (3), gives our main result for the strong-selection regime:

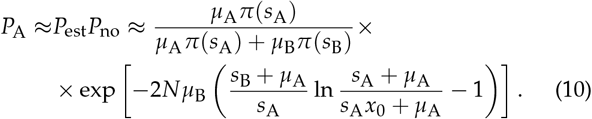

When the population size *N* is small, *N*_*e*_ is correspondingly small as well, so that *P*_no_ is close to 1. In this limit, Eq. (10) reduces to the simpler SSWM approximation *P*_A_ *≈P*_est_. We therefore expect Eq. (10) to remain a good approximation even when the population is small. In Fig. 3, we compare Eq. (10) to results from the direct simulation of the full model and find excellent agreement. More comparison plots can be found in SI Fig. 1. We note that, in order to obtain good agreement even for small *N*, we use the accurate expression (2) for *π*(*s*) in the evaluation of *P*_est_ and *x*_0_ rather than the approximation *π*(*s*) *≈*2*s*.

**Figure 3.**
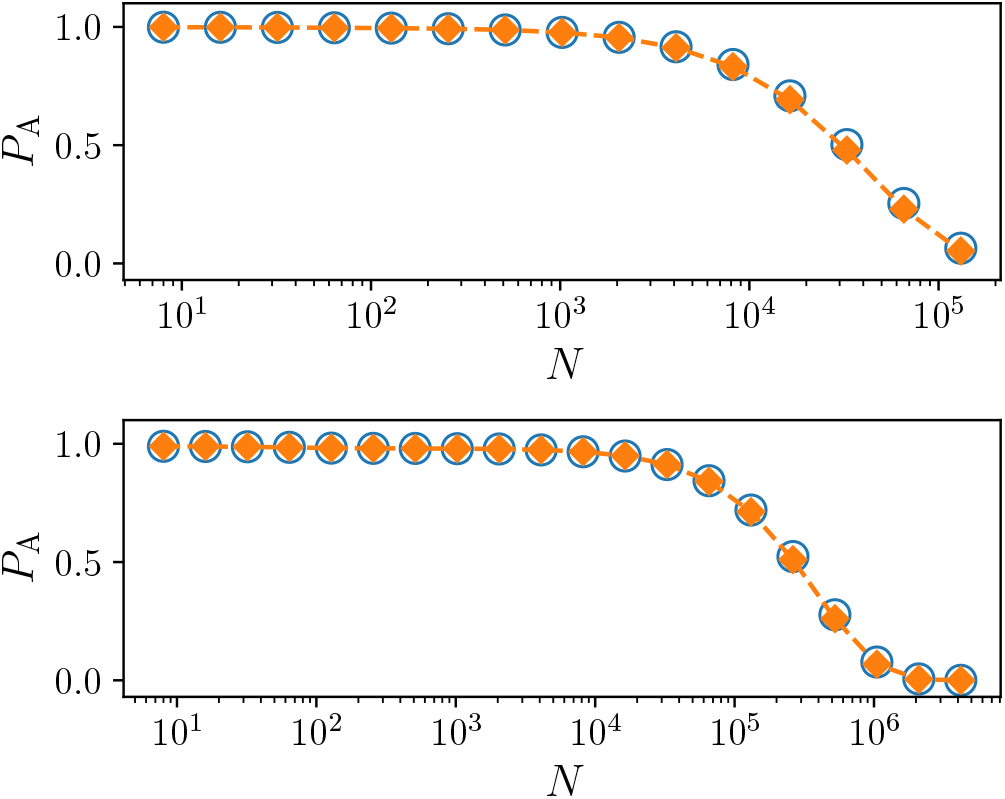
Plots of *P*_A_ vs. *N* on a semi-logarithmic scale. The parameters are (top) *s*_A_ = 0.01, *s*_B_ = 0.05, *µ*_A_ = 10^*−*3^, *µ*_B_ = 10^*−*6^ and (bottom) *s*_A_ = 0.01, *s*_B_ = 0.02, *µ*_A_ = 10^*−*5^, *µ*_B_ = 10^*−*7^. In every panel, the empty circles present simulation results and the filled diamonds show the numerical evaluation of Eq. (10).

Following Yampolsky and Stoltzfus (2001), we also examined the resulting bias in outcomes, defined as the ratio *P*_A_/*P*_B_ = *P*_A_/(1*− P*_A_) of the two fixation probabilities. In Fig. 4, we plot this bias in outcomes as a function of the mutation bias *B ≡ µ*_A_/*µ*_B_, comparing values obtained from direct simulation with the prediction of Eq. (10), and again find excellent agreement. As reported by Yampolsky and Stoltzfus (2001), the bias in outcomes increases linearly with *B* once *B* becomes sufficiently large, but the slope of this linear relation decreases with increasing *N*. To explain this observation, we examine the limit *B→* ∞ by taking *µ*_B_ *→*0 at fixed *µ*_A_. The expression (10) is then of the form

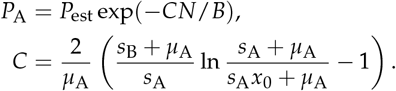

**Figure 4.**
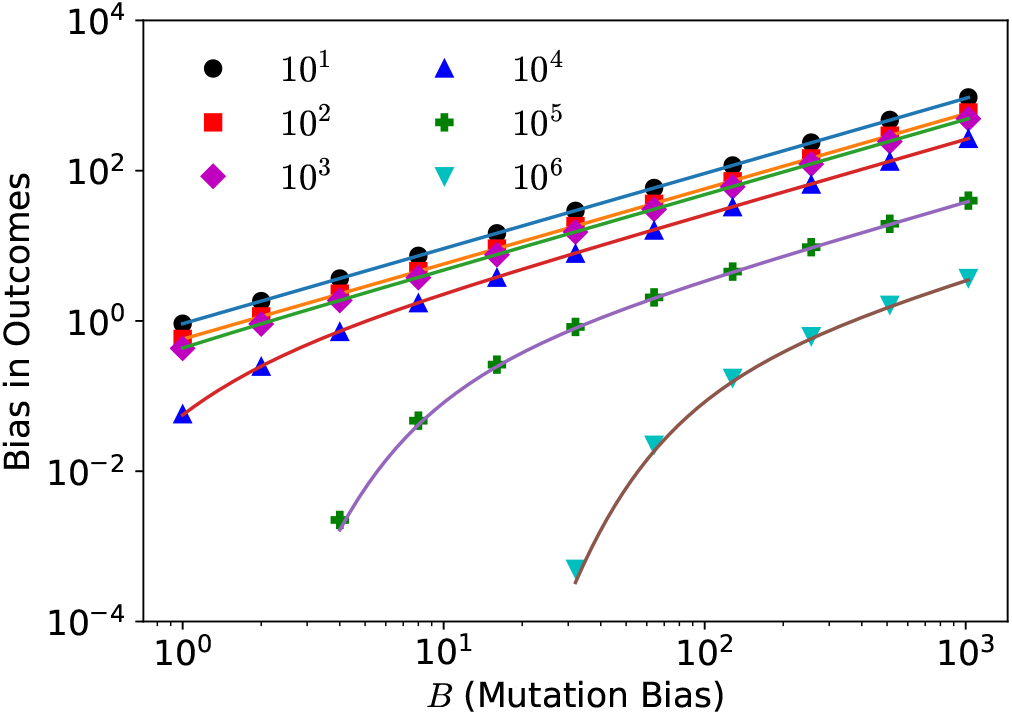
Bias in outcomes *P*_A_/*P*_B_ vs mutation bias *B* = *µ*_A_/*µ*_B_ for *N* = 10^1^, 10^2^, … , 10^6^ (top to bottom) on a double logarithmic scale. Symbols are from simulations and solid curves are from theory. The parameters are *s*_A_ = 0.01, *s*_B_ = 0.02, *µ*_A_ = 10^*−*5^. The theory predicts the bias in outcomes with high accuracy. Note the linear dependence on the mutation bias for large *B*.

For large *B, B ≫ CN*, this implies

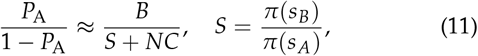

where *S* becomes the selection bias, *s*_*B*_ /*s*_*A*_, when *π*(*s*) *≈*2*s*. The bias in outcomes remains linear in the mutation bias *B*, but the proportionality constant decreases when *NC* become of order unity. The latter condition is easily checked to coincide with the condition for the onset of clonal interference for A mutants, *µ*_*A*_ *N* ln *N >* 1 (Park *et al*. 2010).

In the above study, we have only considered the model with discrete, non-overlapping generations. It is natural to ask how the fixation probability *P*_A_ changes for models with overlapping generations. In Appendix B, we extend our results to the Moran model. The theory remains essentially unchanged, except that *π*(*s*) is replaced by the corresponding expression for the Moran model. There, we also provide a model-independent formula for *P*_A_; see equations (B.4).

### Weak selection

By symmetry, it is immediately clear that when *s*_A_ = *s*_B_, we must have *P*_A_ = *µ*_A_/(*µ*_A_ + *µ*_B_), regardless of how large *N* is. However, taking the limit *s*_B_ *→s*_A_ in Eq. (10) fails to reproduce this obvious result. This discrepancy arises because Eq. (10) treats A and B asymmetrically, effectively assuming that B is destined to fix as soon as it becomes established. That assumption breaks down precisely when *s*_A_*≈ s*_B_ (i.e., under weak selection between the two mutants), since in that case A retains a non-negligible chance of fixing even after B has already become established. Equation (10) is therefore not applicable in the weak-selection regime, and a different approach is needed, which we now develop.

When *s*_A_*≈ s*_B_, we no longer need to distinguish between A and B up to the point at which the wild-type genotype disappears from the population entirely. Neglecting the small difference between the two selection coefficients, the frequency *f* _*A*_ of A at the moment the wild type goes extinct is determined by the balance of the mutation processes as

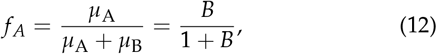

where, as before, *B* = *µ*_A_/*µ*_B_ denotes the mutation bias. At this point no further new mutations need be considered, so we can apply Kimura’s fixation probability formula generalized to the case where A is initially present at frequency *f* _*A*_ (Kimura 1962). This yields

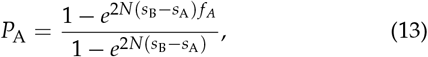

which correctly approaches *f* _*A*_ in the limit *s*_B_*→ s*_A_, as required. In Fig. 5, we compare simulation results for weak selection with Eqs. (13) and (10). Since Δ*s* is very small, the strong selection formula obviously fails to predict the simulation results, while the weak selection formula gives a very accurate prediction. Results for additional parameter combinations are presented in SI Fig. 2. As Δ*s* := *s*_B_*− s*_A_ diminishes, Eq. (13) becomes progressively more accurate, while Eq. (10) becomes progressively worse, eventually failing outright as we anticipated before. It might seem tempting to conclude simply that Eq. (10) is the right choice when *N*Δ*s* is large and Eq. (13) is the right choice when *N*Δ*s* is small. This criterion, however, is not exclusive: there is an intermediate regime in which both equations give good agreement with simulation, as shown in the bottom panel of Fig. 6. Results for additional parameter combinations are presented in SI Fig. 3, which shows how progressively agreement is achieved.

**Figure 5.**
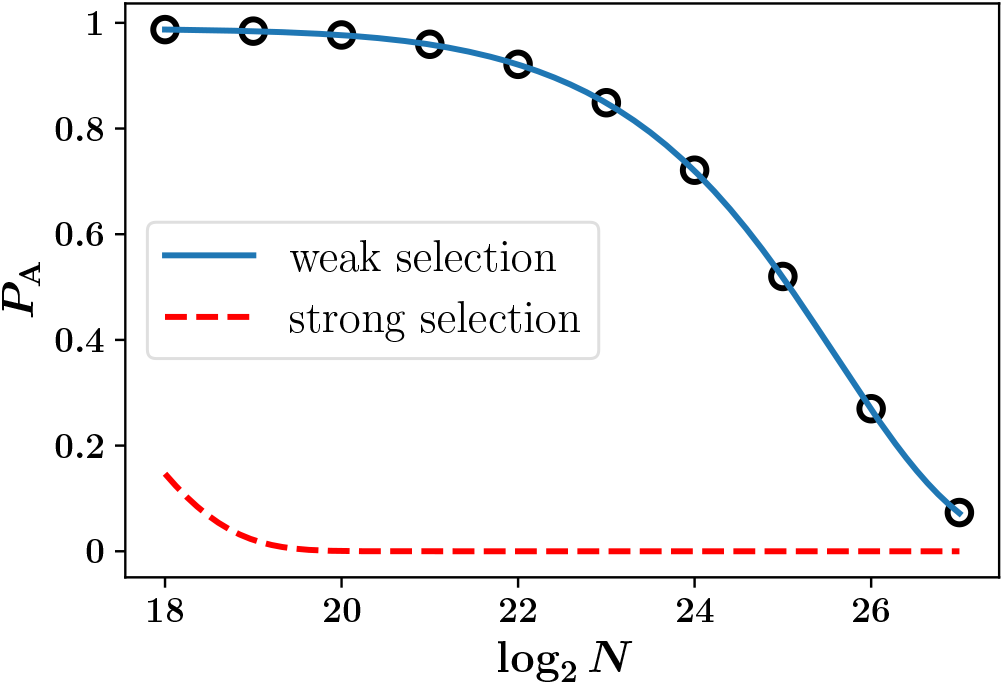
Plots of *P*_A_ vs. log_2_ *N* for *s*_A_ = 10^*−*2^, Δ*s* = 10^*−*6^, *µ*_A_ = 10^*−*4^, and *µ*_B_ = 10^*−*6^. The weak selection formula (13) (solid blue curve) and the strong selection formula (10) (red dashed curve) are compared.

**Figure 6.**
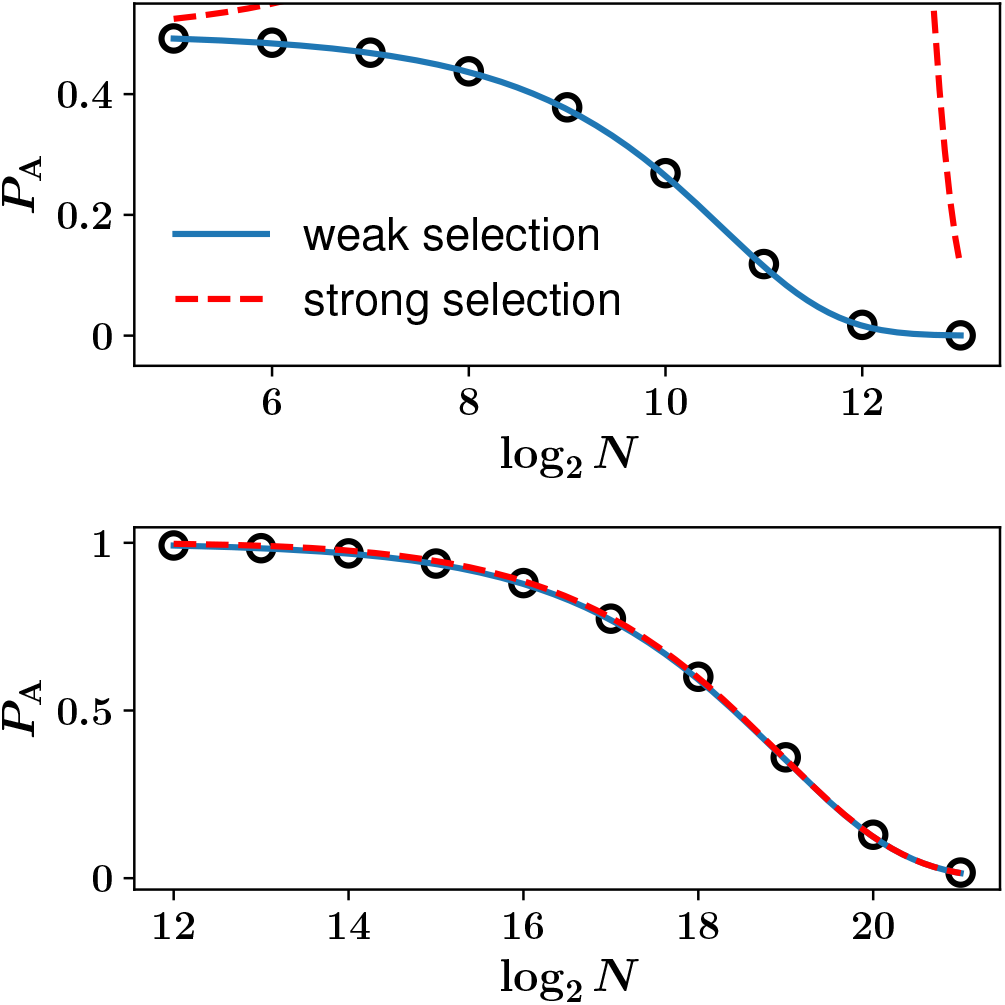
Plots of *P*_A_ vs log_2_ *N* for *s*_A_ = 10^*−*4^, Δ*s* = 10^*−*3^, *µ*_A_ = 10^*−*3^, and (top) *µ*_B_ = 10^*−*3^, (bottom) *µ*_B_ = 10^*−*6^. Circles present simulations results. The weak selection formula (solid blue curve) performes better than the strong selection formula (red dashed curve) in general for these parameter sets, but for *µ*_B_ = 10^*−*6^ (bottom) both formulae are accurate.

Interestingly, the bias in outcomes is linear in the mutation bias for large *B* also in the weak selection regime. Noting that 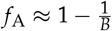 for large *B* and evaluating (13) in this limit yields

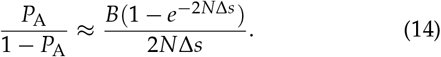

In the neutral limit, Δ*s→* 0, the bias in outcome is equal to the mutation bias, but for Δ*s >* 0 the prefactor of the linear relation is reduced, similar to (11). Note that (13) and (14) hold irrespective of the sign of Δ*s*, and can be applied also when mutation bias and selection are aligned, in the sense that *µ*_*A*_ *> µ*_*B*_ and *s*_A_ *> s*_B_, a situation observed experimentally by Barber and Couce (2026). In this case (14) shows, rather obviously, that the bias in outcomes exceeds the mutation bias.

## Discussion

### Summary and relation to previous work

We have derived closed-form expressions for the fixation probability *P*_A_ of the YS model across the previously unresolved intermediate regime of appreciable mutation supply, addressing the gap identified by Stoltzfus (2019). We obtained Eq. (10) for the strong-selection, that is, ECI framework and Eq. (13) for the weak-selection (near-neutral) framework. Our resulting formulas are found to be in good agreement with simulation results of the full model in their respective regime of validity, and together they cover the great majority of biologically relevant combinations of *N, µ*_A_, *µ*_B_, *s*_A_, and *s*_B_. Both expressions also recover the known limiting cases. Equation (10) reduces to the SSWM/origin-fixation prediction *P*_A_ *≈P*_est_ when *N* is small, and Eq. (13) reduces to the purely mutation-driven result *P*_A_ = *µ*_A_/(*µ*_A_ + *µ*_B_) as *s*_B_*→ s*_A_. In this sense, the ECI framework introduced here can be viewed as a generalization of the SSWM/origin-fixation framework rather than a replacement for it.

The core idea behind the ECI approach was already introduced by Jain *et al*. (2011). We introduce two important corrections relative to their earlier treatment. The first is to retain the *µ*_A_ term describing the ongoing effect of mutation during the growth phase. The second is to allow the establishment probability of fitter mutant in the growth phase to depend on the actual frequency of *A*, rather than treating it as a constant. As shown by Jain *et al*. (2011), neglecting these two factors still gives reasonably accurate results as long as *µ*_A_/*µ*_B_ is not too large, but the result becomes progressively less accurate as this ratio grows, that is, in the regime of strong mutation bias that motivates this entire line of research.

### Resolving the controversy

Our results bear on the recent debate over whether mutation-biased adaptation persists in large populations. Svensson and Berger (2019) have argued that the effect should weaken as population size grows, while Barber and Couce (2026) report experimental evidence that it need not, and can even intensify, depending on the nature of the bias. Equation (10) offers a route to resolving this issue. At heart, the controversy arises from the non-exchangeability of the limits of large population size and large mutation bias. Taking the limit *N →*∞ at fixed mutation rates and selection coefficients, the probability *P*_A_ for mutant A to fix vanishes, indicating that selection dominates the process. On the other hand, in the limit of large *B* at fixed population size, the bias in outcomes remains proportional to the mutation bias, but with a constant of proportionality that is a decreasing function of *N*, see (11) and also (14).

Our results thus clarify, in the context of the simple YS model, the widespread notion that the effect of mutation bias on adaptation weakens with increasing mutation supply (Svensson and Berger 2019; Gomez et al. 2020; Cano et al. 2022, 2023). Cano *et al*. (2022) quantified this effect through a power-law relationship

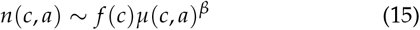

between the observed number of substitutions *n*(*c, a*) from codon *c* to codon *a*, the frequency of codon *c*, and the corresponding mutation rate *µ*(*c, a*), and showed through simulations that the exponent *β* decreases from its SSWM value *β* = 1 towards *β* = 0 as the mutation supply *Nµ* increases. In contrast, for the YS model, the exponent is found to remain at *β* = 1, while the factor of proportionality in the relation (15) (which is represented in the analysis of Cano *et al*. (2022) by the intercept *β*_0_ in the logarithmic regression between *n* and *µ*) decreases.

### Application to a microbial evolution experiment

To illustrate the applicability of our result to a specific experimental system, we revisit the work of Schenk *et al*. (2022) on the adaptation of small (*N* = 2 *×*10^6^) and large (*N* = 2 *×*10^8^) populations of *E. coli* to increasing levels of the *β*-lactam antibiotic cefotaxime. After evolving replicate populations over 500 generations, it was observed that adaptation in large populations was driven by single nucleotide polymorphisms (SNPs), whereas small populations preferentially adapted by fixing large and intermediate scale structural variants (insertions and deletions). It was hypothesized that these alternate routes reflected a tradeoff between rate and effect size for the two classes of substitutions, where structural variants of high rate and small effect are favored in small populations, whereas SNPs of low rate and large effect dominate in large populations. A computational analysis of the mutational content of the endpoint clones supported this hypothesis and provided estimates for the mutation rates and selection coefficients for the two mutation classes. In Fig. 7, we evaluate the fixation probability (10) for a pair of mutations, one from each class, with the inferred parameters. While this comparison is oversimplified in many respects, our theory predicts the transition from adaptation through high rate/low effect mutations to low rate/high effect mutations in the range of population sizes considered in the experiment.

**Figure 7.**
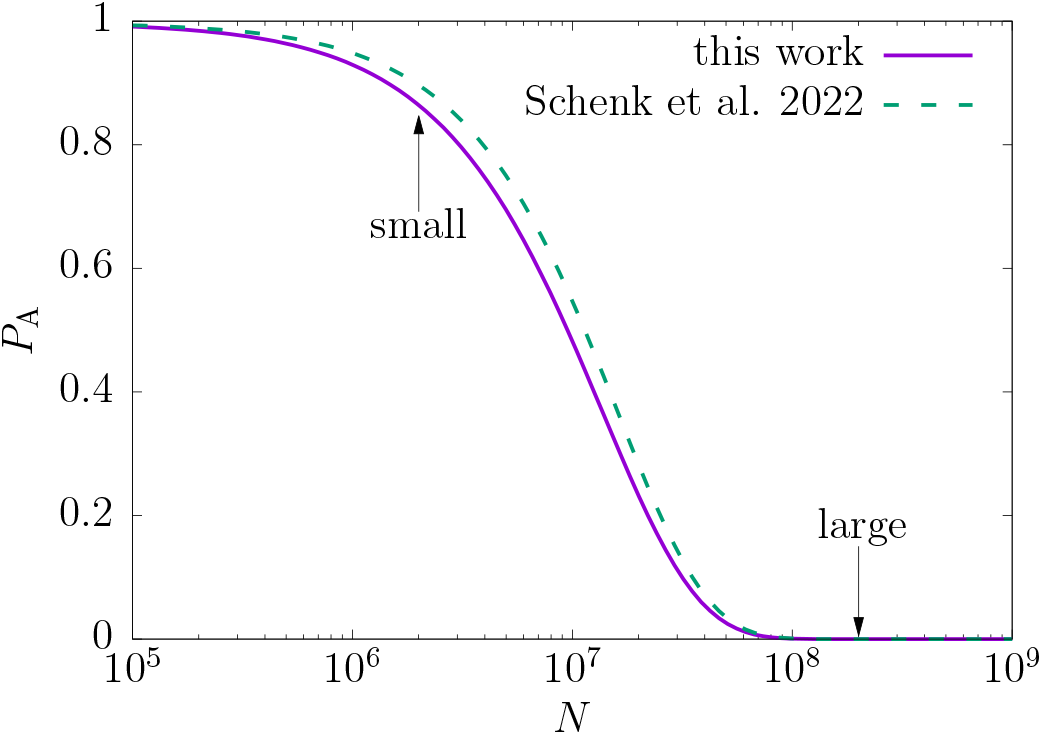
Plot of *P*_A_ vs *N* for selection coefficients and mutations rates inferred by Schenk *et al*. (2022) for two classes of mutations arising in small and large populations of *E. coli* adapting to increasing concentrations of the *β*-lactam antibiotic cefotaxime. Specifically, selection coefficient *s*_A_ = 0.14 and *s*_B_ = 0.41 were inferred computationally for large-scale structural variants (type A) and single-nucleotide polymorphisms (type B), respectively, and the corresponding mutation rates for single mutations in each class were *µ*_A_ = 1.2*×* 10^*−*6^ and *µ*_B_ = 1.1*×* 10^*−*9^. An approximate *ad hoc* formula for *P*_A_ proposed by Schenk *et al*. (2022) is plotted for comparison (dashed curve).

### Outlook

We have considered only two competing mutational routes, whereas adaptation in real populations, for example, resistance evolution against a given antibiotic, often proceeds via a larger set of *K >* 2 mutationally accessible routes competing simultaneously (Orr 2005; Schenk *et al*. 2012; Das and Krug 2022; Schenk *et al*. 2022). The establishment-clonal interference (ECI) approach generalizes naturally to this situation when the quantity of interest is the fixation probability of the least fit genotype A. The same logic developed for the two-route case applies directly. First, the genotype A must become established before any other genotypes, since otherwise its probability of eventual fixation is almost zero. Second, no fitter genotype may become established during the growth phase of A. Under these conditions the calculation extends straightforwardly, and is detailed in Appendix C; see equations (C.2).

In contrast, an explicit formula for fixation probability of, say, the *i*-th genotype is difficult, if not impossible, to obtain. The difficulty stems from having to also deal with the probability that the *i*-th genotype becomes established during the growth phase of a less fit genotype, along with the constraint that no fitter genotype should be established during the growth phase of the *i*-th genotype. This difficulty can be circumvented only in the two-route case, because *P*_B_ = 1 *−P*_A_ by construction.

Extending the weak selection scenario to *K >* 2 genotypes requires that *N* |*s*_*i*_*− s*_*j*_| *≪*1 for all pairs *i, j*. Just as in Section Weak selection, the difference in selection coefficients can be ignored until the wild type goes extinct. Subsequently, the problem reduces to finding the fixation probabilities of the different variants from standing variation with initial frequencies *x*_*i*_ = *µ*_*i*_ / Σ_*j*_ *µ*_*j*_, *i* = 1, , *K*. This problem was studied, e.g., by Overall and Waxman (2024), but a full analytic expression does not seem to be available. We leave this question for future work. Another natural extension is to include the effect of recombination. Here, epistasis must be taken into account as a key factor. If reciprocal sign epistasis is present, that is, if the double mutant carrying both A and B has lower fitness than either single mutant, recombination between the A and B backgrounds offers little advantage, since the recombinant genotype carrying both mutations is disfavored by selection. In this case recombination can effectively be neglected, and the dynamics should reduce to those already captured by the ECI framework developed above. When epistasis is of magnitude type or absent instead, recombination can bring the two beneficial mutations together onto a single background, which fundamentally changes the competition among routes and requires a separate treatment.

Each of these extensions would relax one of the simplifying assumptions discussed above, moving the theory progressively closer to the complexity of real adaptive landscapes while retaining the analytical tractability that has been the central aim of this work.

## Data availability

All data generated and analyzed during this study are included in this publication.

## AI Statement

LLM tools (Claude) were used as support for language editing and literature search.

## Appendix A

Master equation for the Yampolsky-Stoltzfus model and deterministic dynamics in the growth phase

In this appendix we derive Eq. (4) from the master equation for the Wright-Fisher sampling dynamics. We first write down the master equation. Let *N*_0_, *N*_*A*_, and *N*_*B*_ denote the random variables for the numbers of wild-type, mutant A, and mutant B individuals, respectively. We assume a fixed population size *N* across all generations, so that *N*_0_ + *N*_*A*_ + *N*_*B*_ = *N*. Conditioned on *N*_*i*_ = *n*_*i*_ at generation *t*, the probability that *N*_*i*_ = *m*_*i*_ at generation *t* + 1 (*i ∈ {*0, *A, B}*) is given by the multinomial (Wright–Fisher) sampling rule

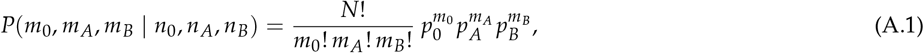

where

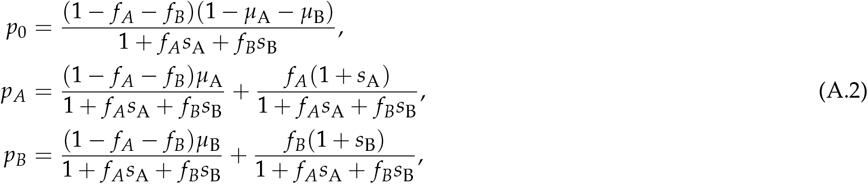

with *f* _*A*_ = *n*_*A*_ /*N* and *f*_*B*_ = *n*_*B*_ /*N* the frequencies of mutant A and mutant B, so that 1 *−f* _*A*_ *−f*_*B*_ = *n*_0_/*N* is the wild-type frequency, and the relative fitness advantages defined as

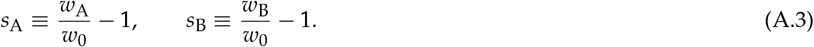

Writing *P*(*n*_0_, *n*_*A*_, *n*_*B*_; *t*) for the probability that *N*_*i*_ = *n*_*i*_ at generation *t* for *i ∈ {*0, *A, B}*, the master equation follows as

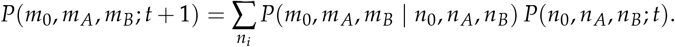

We now derive Eq. (4). Since we are only interested in the growth of the mutant A frequency, we set *f*_*B*_ = *µ*_B_ = 0 in equations (A.1), effectively reducing the dynamics to wild type and mutant A alone. Let *P*(*m, t*) denote the probability of observing *m* copies of mutant A at generation *t*. The dynamical rule then relates *P*(*n, t* + 1) to *P*(*m, t*) via

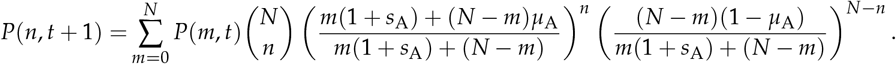

Using 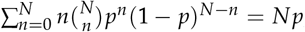, the mean number of mutants at generation *t* + 1 satisfies

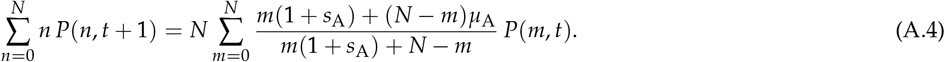

Denoting the average frequency of mutant A at generation *t* by 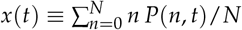, and neglecting fluctuations on the right-hand side of equations (A.4), we obtain

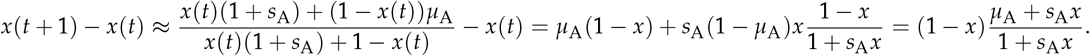

Interpreting *x*(*t* + 1) *− x*(*t*) as a time derivative, we arrive at Eq. (4).

## Appendix B: Extension to the Moran model and other reproduction rules

The results in the main text were derived under the non-overlapping-generation Wright-Fisher model. Many biological populations of interest, including the bacterial systems used in recent experimental tests of mutation-biased adaptation (Barber and Couce 2026), however, are better described by overlapping generations and continuous-time reproduction, as captured by the Moran model. Extending the present analysis to the Moran model is quite straightforward.

In the Moran model, it is convenient to define the mutant selection coefficients in terms of the mutant and WT fitnesses as *s*_*i*_ = 1 *− w*_0_/*w*_*i*_ , *i* = A,B. The Kimura formula (2) for the fixation probability can then be replaced by the exact expression

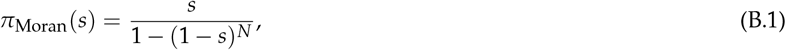

which reduces to *π*(*s*) *≈ s* for *s >* 0 and large *N*. With this modification, the analysis in Section Results is applicable almost verbatim. Setting *π*_Moran_(*s*_B_ *− s*_A_ *x*) = *s*_B_ *− s*_A_ *x* in Eq. (5), we get

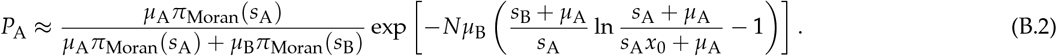

We performed simulations of the Moran model, finding good agreement with equations (B.2); see SI Fig. 4.

Note that the factor in front of *Nµ*_B_ in the exponent of (10) and (B.2) is the proportionality constant *α* in the relation *π*(*s*) *≈αs* for small *s*. A linear dependence of the fixation probability on the selection coefficient for large *N* and small *s* holds for other reproduction rules as well, see e.g. Johnson and Gerrish (2002) for the case of binary fission, and we can exploit this observation to derive a universal form of *P*_A_. Using *x*_0_ = 1/(*Nπ*(*s*_A_)) = 1/(*αNs*_A_) and defining (*i ∈ {*A, B*}*)

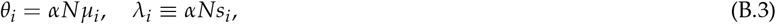

we obtain a model-independent formula

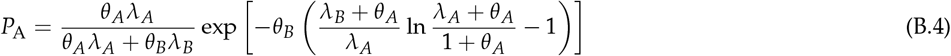

that reduces to (10) and (B.2) for *α* = 2 and *α* = 1, respectively. If we consider a random mating diploid population of size *N* without dominance, we expect that equations (B.4) remains applicable by taking *N 1→* 2*N*.

## Appendix C: Fixation probability of least favorable genotype among *K* beneficial mutations

In this appendix, we derive the fixation probability for the selectionally least favored path among *K* mutational paths. To be specific, we denote the least favored genotype by A and the other genotypes by *i*, 2*≤ i ≤K*. The mutation rate and selection coefficient of the *i*-th genotype are denoted by *µ*_*i*_ and *s*_*i*_ , respectively. As summarized in Section Outlook, the fixation probability *P*_A_ is the product of the establishment probability *P*_est_ and the probability *P*_no_ that no fitter genotype than A can establish during the growth phase.

We first calculate *P*_est_, generalizing Eq. (3) to this case. Since A should be established before the establishment of other genotypes, we have

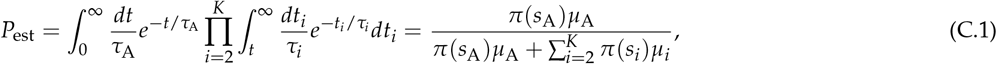

where *τ*_*i*_ is as given in Eq. (1). In the growth phase, the expected number of establishment events of genotype *i* is obtained by replacing B by *i* in Eq. (7). Therefore, the desired fixation probability is

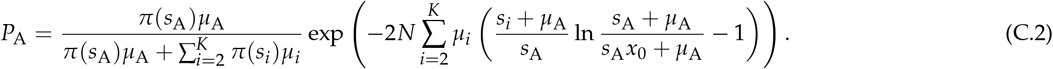

This expression reduces, as expected, to the result by McCandlish and Stoltzfus (2014) in the SSWM/origin-fixation regime, *Nµ*_*i*_ ≪ 1. We note that the above fixation probability is valid regardless of the ordering of the mutation rates as long as *s*_A_ is the smallest among the beneficial selection coefficients.

**SI Figure 1.**
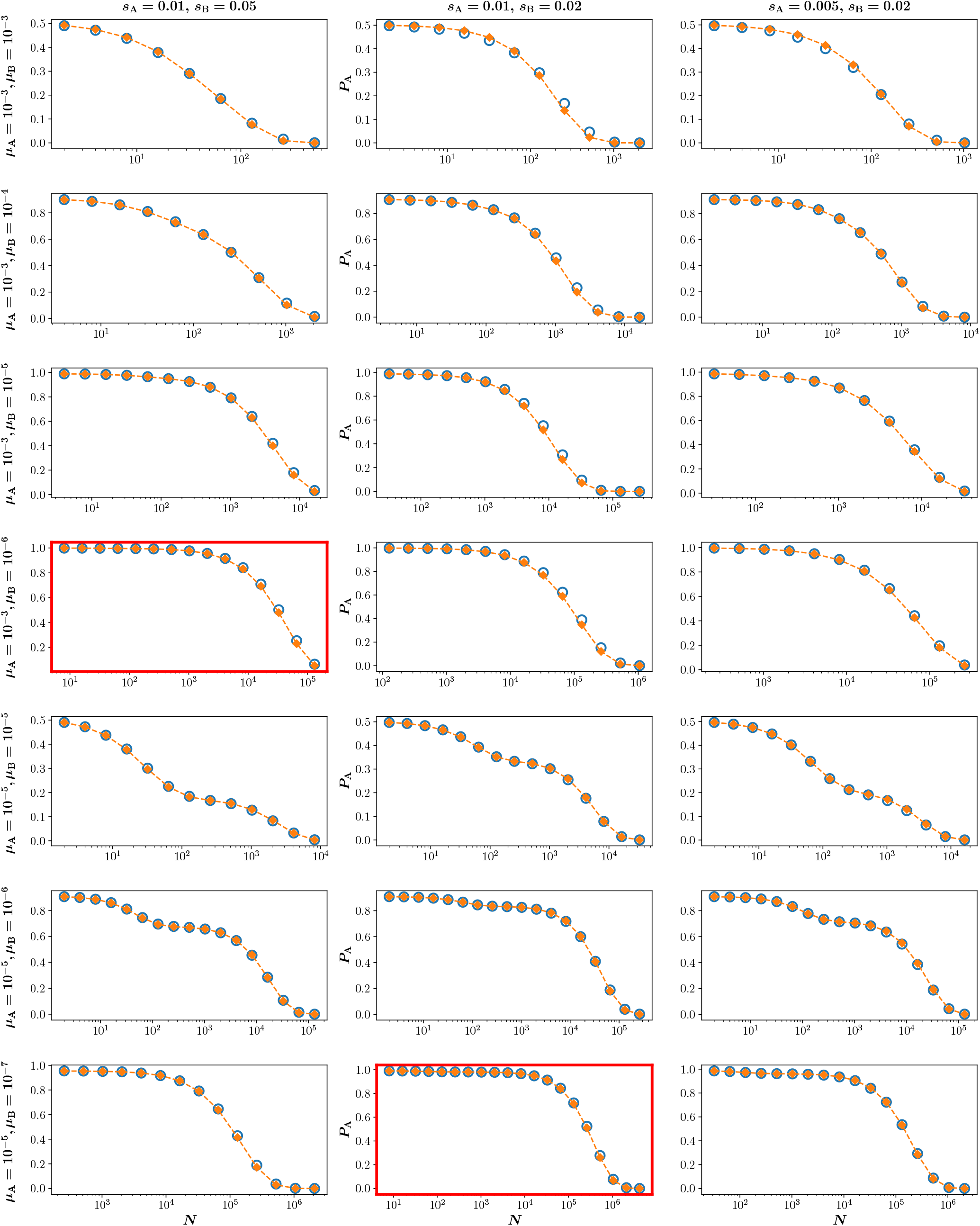
Plots of *P*_A_ vs. *N* for *s*_A_ = 0.01, *s*_B_ = 0.05 (left column), *s*_A_ = 0.01, *s*_B_ = 0.02 (middle column), and *s*_A_ = 0.005, *s*_B_ = 0.02 (right column). Each row shares the mutation rate (*µ*_A_, *µ*_B_; written on the left). In every panel, the empty circles present simulation results and the filled diamond show numerical calculations of Eq. (10). The red-colored panels are shown in Fig. 3 of the main text.

**SI Figure 2.**
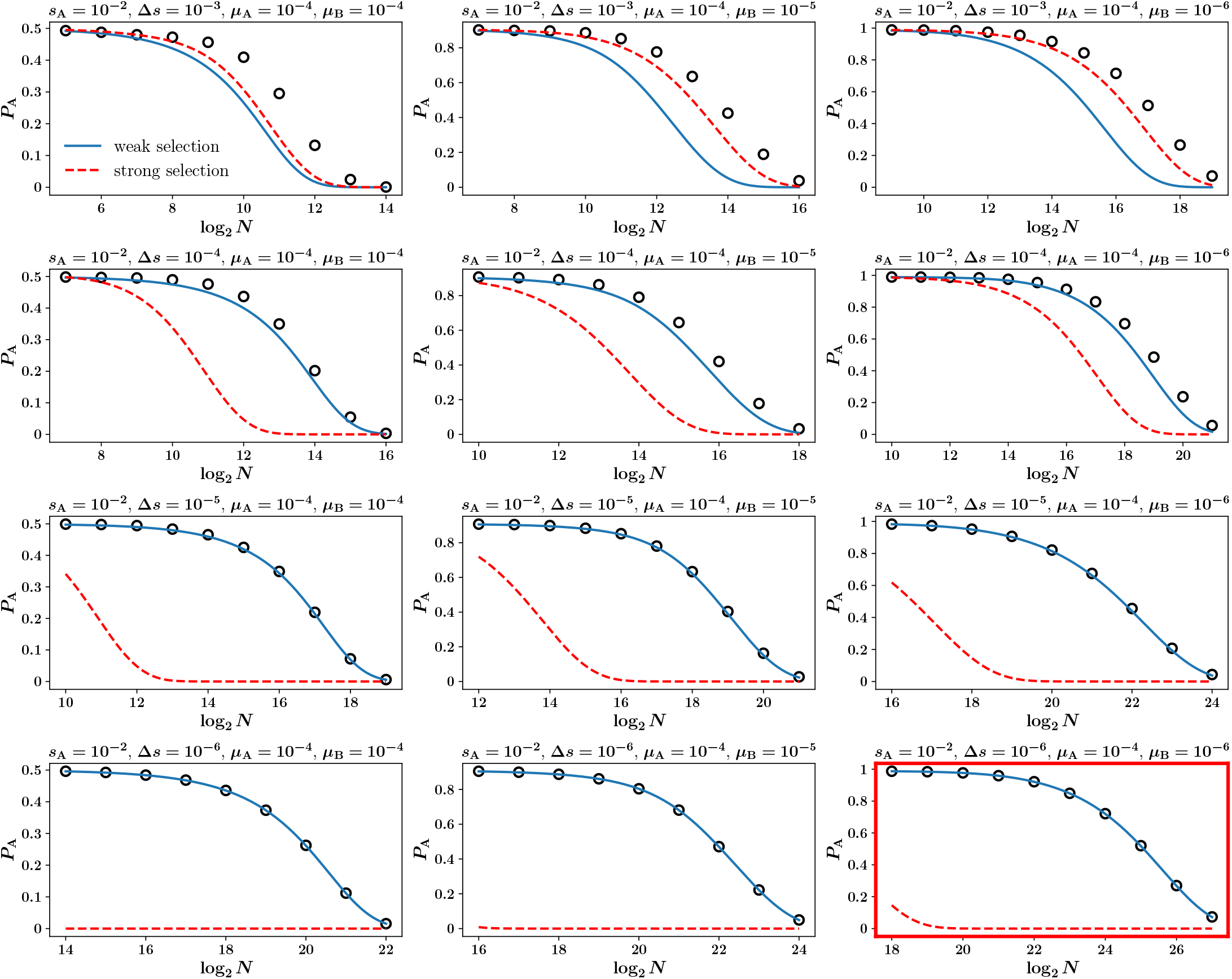
Plots of *P*_A_ vs log_2_ *N* for *s*_A_ = 10^*−*2^ and *µ*_A_ = 10^*−*4^. The values of Δ*s* are 10^*−*3^, 10^*−*4^, 10^*−*5^, and 10^*−*6^ (top to bottom). The values of *µ*_B_ are 10^*−*4^, 10^*−*5^, and 10^*−*6^ (left to right). Weak selection formula (solid blue curve) and strong selection formula (red dashed curve) are compared. The red-colored panel is shown in Fig. 5 of the main text.

**SI Figure 3.**
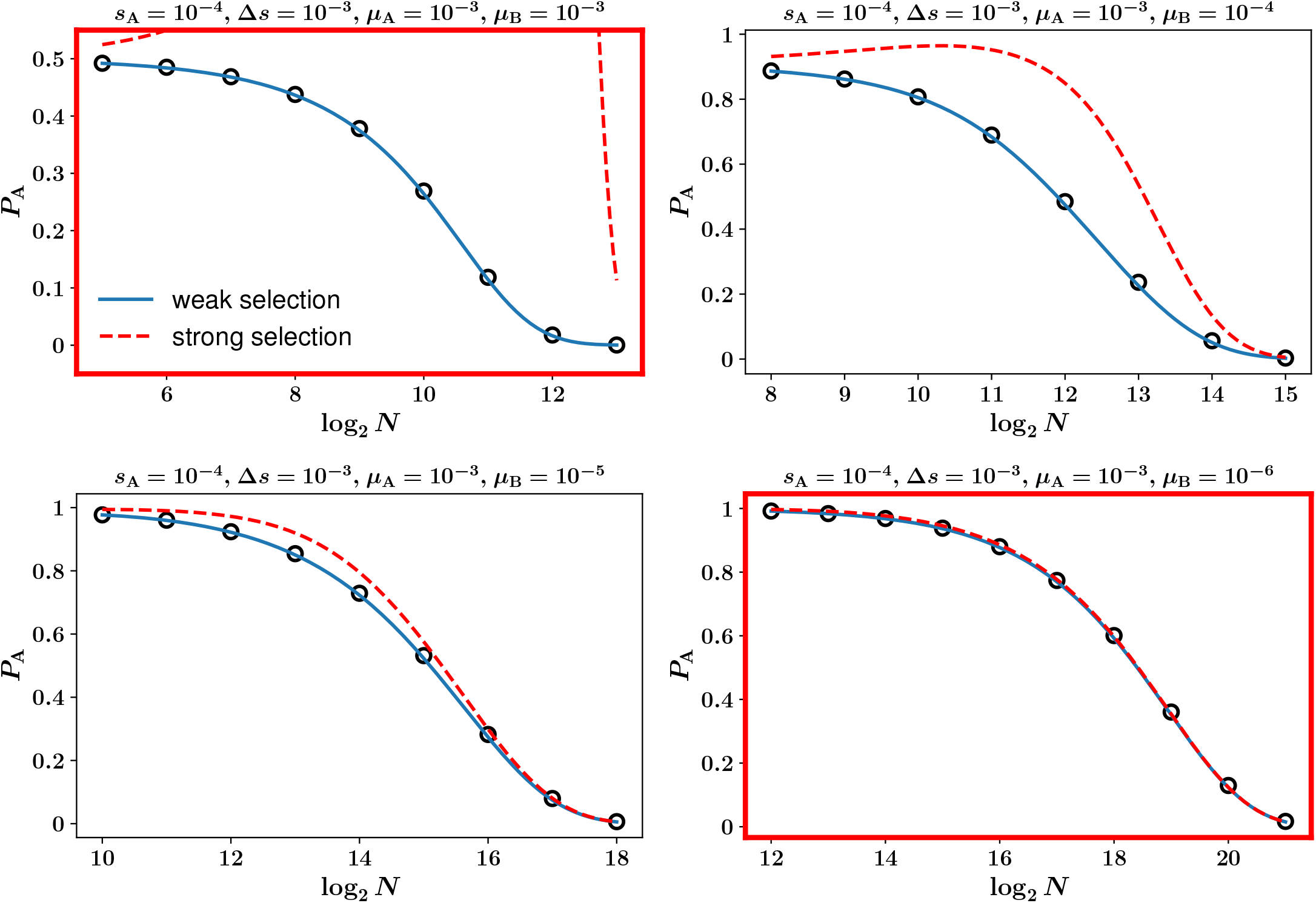
Plots of *P*_A_ vs log_2_ *N*. The values of parameters are given on top of each panel. The red-colored panels are shown in Fig. 6 of the main text.

**SI Figure 4.**
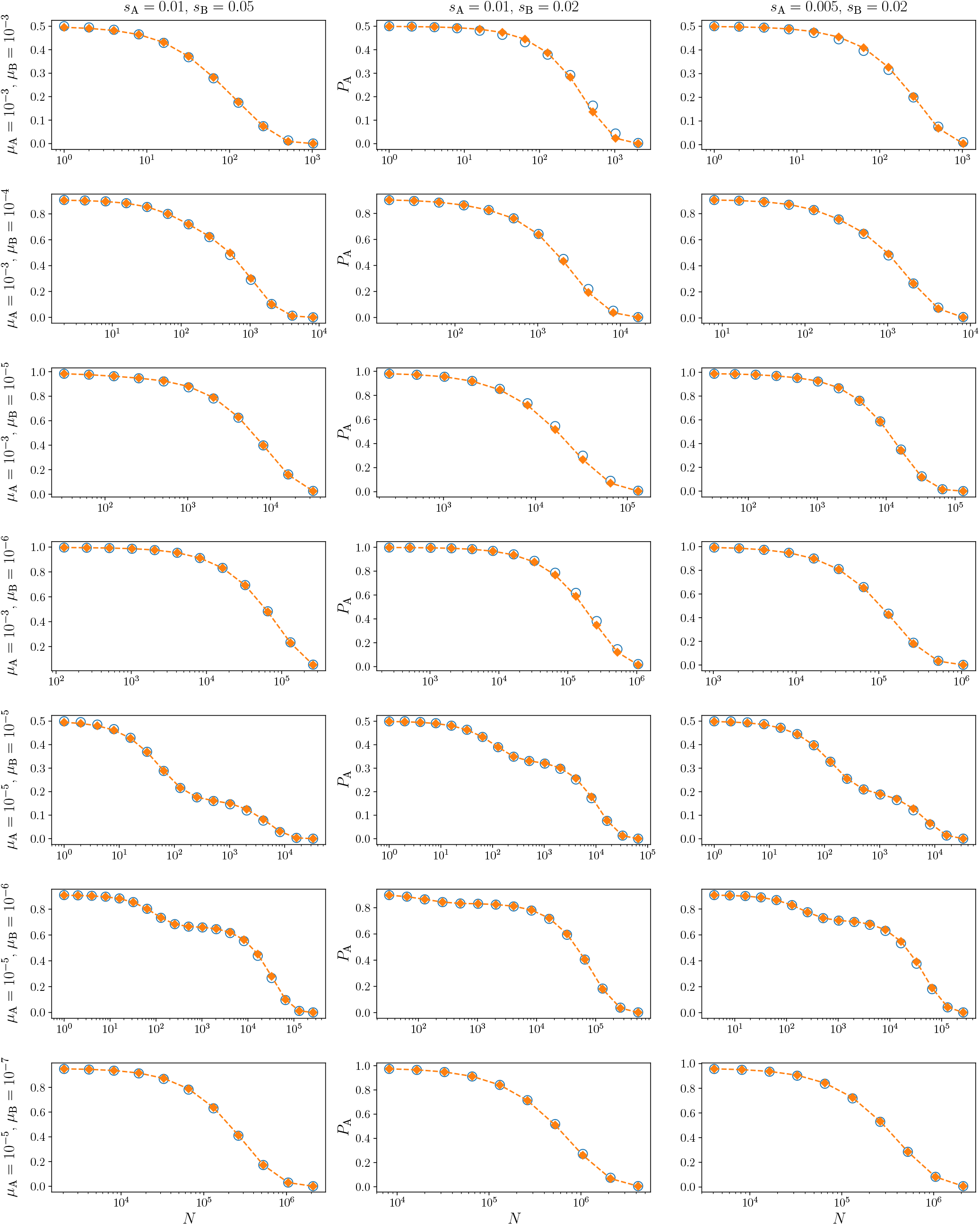
Semi-logarithmic plots of *P*_A_ vs. *N* for the Moran model. Selection coefficients are *s*_A_ = 0.01, *s*_B_ = 0.05 (left column), *s*_A_ = 0.01, *s*_B_ = 0.02 (middle column), and *s*_A_ = 0.005, *s*_B_ = 0.02 (right column). Each row shares the mutation rate (*µ*_A_, *µ*_B_; written on the left). In every panel, the empty circles present simulation results and the filled diamond show numerical calculations of equations (B.2).

## Literature cited

Agashe D, Sane M, Singhal S. 2023. Revisiting the role of genetic variation in adaptation. Am. Nat.. 202:486–502.

Barber JN, Couce A. 2026. Mutation-biased adaptation is consequential even in large bacterial populations. Nat. Comm.. 17:7357.

Cano AV, Gitschlag BL, Rozhoñová H, Stoltzfus A, McCandlish DM et al. 2023. Mutation bias and the predictability of evolution. Philos. Trans. R. Soc. B. 378:20220055.

Cano AV, Rozhoñová H, Stoltzfus A, McCandlish DM, Payne JL. 2022. Mutation bias shapes the spectrum of adaptive substitutions. Proc. Natl. Acad. Sci. USA. 119:e2119720119.

Das SG, Krug J. 2022. Unpredictable repeatability in molecular evolution. Proceedings of the National Academy of Sciences. 119:e2209373119.

Gerrish PJ, Lenski RE. 1998. The fate of competing beneficial mutations in an asexual population. Genetica. 102-103:127– 144.

Gitschlag BL, Stoltzfus A, McCandlish DM. 2025. Molecular adaptation reflects taxon-specific mutational biases. bioRxiv. Preprint.

Gomez K, Bertram J, Masel J. 2020. Mutation bias can shape adaptation in large asexual populations experiencing clonal interference. Proc. R. Soc. B. 287:20201503.

Jain K, Park SC, Krug J. 2011. Evolutionary advantage of small populations on complex fitness landscapes. Evolution. 65:1945– 1955.

Johnson T, Gerrish PJ. 2002. The fixation probability of a beneficial allele in a population dividing by binary fission. Genetica. 115:283–287.

Katju V, Bergthorsson U. 2019. Old trade, new tricks: insights into the phenotypic, population genetic, and molecular consequences of mutations from mutation accumulation experiments. Genome Biol. Evol.. 11:136–165.

Kimura M. 1962. On the probability of fixation of mutant genes in a population. Genetics. 47:713–719.

McCandlish DM, Epstein CL, Plotkin JB. 2015. Formal properties of the probability of fixation: identities, inequalities and approximations. Theor. Popul. Biol.. 99:98–113.

McCandlish DM, Stoltzfus A. 2014. Modeling evolution using the probability of fixation: history and implications. Q. Rev. Biol.. 89:225–252.

Ohta T. 1992. The nearly neutral theory of molecular evolution. Annu. Rev. Ecol. Syst.. 23:263–286.

Orr HA. 2005. The probability of parallel evolution. Evolution. 59:216–220.

Overall A, Waxman D. 2024. Influence of selection on the probability of fixation at a locus with multiple alleles. BMC Genomics. 25:819.

Park SC, Krug J. 2007. Clonal interference in large populations. Proc. Natl. Acad. Sci. USA. 104:18135–18140.

Park SC, Simon D, Krug J. 2010. The speed of evolution in large asexual populations. J. Stat. Phys.. 138:381–410.

Poelwijk FJ, Kiviet DJ, Weinreich DM, Tans SJ. 2007. Empirical fitness landscapes reveal accessible evolutionary paths. Nature. 445:383–386.

Salverda ML, Dellus E, Gorter FA, Debets AJ, Van Der Oost J et al. 2011. Initial mutations direct alternative pathways of protein evolution. PLoS genetics. 7:e1001321.

Schenk MF, Szendro IG, Krug J, de Visser JAG. 2012. Quantifying the adaptive potential of an antibiotic resistance enzyme. PLoS genetics. 8:e1002783.

Schenk MF, Zwart MP, Hwang S, Ruelens P, Severing E et al. 2022. Population size mediates the contribution of high-rate and large-benefit mutations to parallel evolution. Nat. Ecol. Evol.. 6:439–447.

Soares AdA, Wardil L, Klaczko LB, Dickman R. 2021. Hidden role of mutations in the evolutionary process. Phys. Rev. E. 104:044413.

Stoltzfus A. 2019. Understanding bias in the introduction of variation as an evolutionary cause, In: Uller T, Lala KN, editors, Evolutionary Causation: Biological and Philosophical Reflections, The MIT Press. Massachusetts.

Stoltzfus A. 2021. Mutation, randomness, and evolution. Oxford University Press. Oxford.

Stoltzfus A, McCandlish DM. 2017. Mutational biases influence parallel adaptation. Mol. Biol. Evol.. 34:2163–2172.

Svensson EI. 2022. Multivariate selection and the making and breaking of mutational pleiotropy. Evol. Ecol.. 36:807–828.

Svensson EI, Berger D. 2019. The role of mutation bias in adaptive evolution. Trends Ecol. Evol.. 34:422–434.

Yampolsky LY, Stoltzfus A. 2001. Bias in the introduction of variation as an orienting factor in evolution. Evol. Dev.. 3:73– 83.

